# Network stress test reveals novel drug potentiators in *Mycobacterium tuberculosis*

**DOI:** 10.1101/429373

**Authors:** Shuyi Ma, Robert Morrison, Samuel J. Hobbs, Jessica Farrow-Johnson, Tige R. Rustad, David R. Sherman

**Affiliations:** Center for Infectious Disease Research, formerly Seattle Biomedical Research Institute, Seattle, Washington, 98109, USA.; Interdisciplinary Program of Pathobiology, Department of Global Health, University of Washington, Seattle, Washington 98195, USA

## Abstract

Deciphering molecular stress response is highly relevant to studies of microbes such as *Mycobacterium tuberculosis* (MTB), the causative pathogen of tuberculosis (TB) which sickens 10 million people and kills 1.8 million each year^1^. Prolonged therapy and unfavorable outcomes arise partially because MTB has evolved stress responses to achieve tolerance, wherein MTB persists in otherwise inhibitory drug concentrations by means independent of heritable resistance mutations^2–4^. Understanding these adaptations and how they are regulated can reveal new biology, including unexplored drug targets and treatment-enhancing strategies. Here, we present a novel network-based genetic screening approach: the Transcriptional Regulator Induced Phenotype (TRIP) screen, which we used to identify previously uncharacterized MTB network adaptations to the first-line drug isoniazid (INH). We found regulators that alter INH susceptibility when induced, several of which could not be identified by standard gene disruption approaches. We then focused on a specific regulator, *mce3R*, which potentiated INH when induced. We compared *mce3R*-regulated genes with the baseline INH transcriptional response and implicated the gene Rv1469 as a putative INH effector. Phenotyping a disruption mutant strain then demonstrated a previously unknown role for Rv1469 in INH susceptibility. Integrating TRIP screening with network data can uncover sophisticated molecular response programs.

Stress response in bacteria has been studied both through targeted characterization of individual genes and by systems-level screens. In-depth characterization of specific genes is necessary to elucidate molecular details, but these efforts are time-and resource-intensive. Systems-level studies often rely on high-throughput phenotypic profiling of transposon mutagenesis libraries (e.g. Tn-seq, TraSH, HITS, IN-seq, TraDIS)^5–12^. These approaches assess differential viability among pools of transposon-mediated gene disruption mutants, enabling unbiased screening for putative candidate genes. While powerful, these methods pose some limitations: 1) they cannot identify genes whose upregulation generates a phenotype, 2) essential genes cannot be profiled, and 3) they miss emergent phenotypes that arise only from coordinated actions of multiple genes.

To address these limitations, we developed a strategy that quantifies growth associated with individually inducing each MTB transcription factor (TF). This <u>T</u>ranscriptional <u>R</u>egulator <u>I</u>nduced <u>P</u>henotype (TRIP) screen exploits a library of 207 TF-induction (TFI) strains, representing 97% of annotated MTB regulators, each transformed with a plasmid carrying an MTB TF under control of a mycobacterial chemically-inducible promoter (see Methods for details) (**Figure 1**). Each strain is engineered for conditional induction of a single MTB TF and expression of its associated regulon—the set of genes whose expression changes when a given TF is induced^13–15^. ChIP-seq and expression profiling of these TFI strains under *in vitro* log-phase conditions revealed a baseline network of transcriptional impacts and DNA binding interactions triggered by each TF^13–15^.

**Figure 1.**
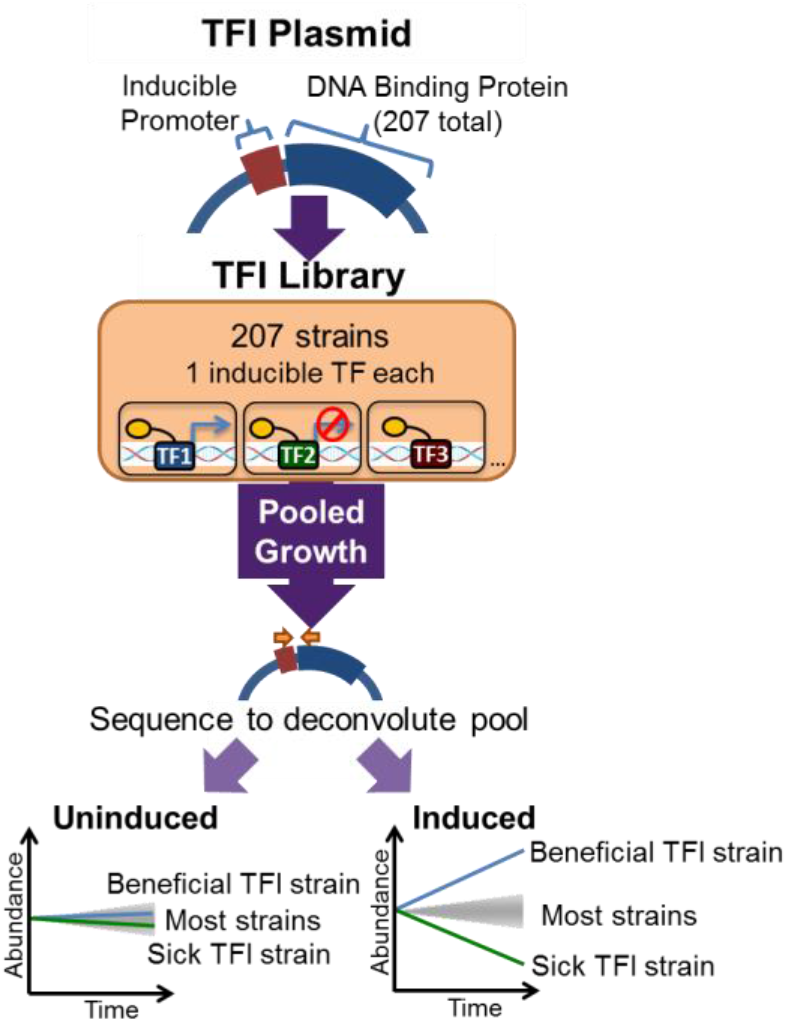
Schematic of TRIP screen.

Here, we pool the TFI strain library for simultaneous measurement of strain growth (**Figure 1**). The pool is exposed to a stress condition either with or without TF induction. The proportion of each TFI strain in the pool is quantified by next-generation sequencing of a DNA segment unique to each strain (see Methods for details). Sampling the pooled culture over time generates simultaneous abundance curves for each TFI strain. The fold abundance change of each strain when induced compared to uninduced identifies regulons with altered growth or survival.

TRIP screens offer several powerful advantages: 1) emergent, network phenotypes are accessible, since regulons generally include multiple genes selected for co-regulation by evolution; 2) revealed phenotypes can be deconstructed with the existing baseline regulatory network; 3) TF expression is chemically triggered, enabling context-specific interrogation of perturbations; and 4) essential regulators and effector genes can be assessed. Moreover, TRIP is broadly extensible—the pipeline can be harnessed to interrogate MTB phenotypes under numerous stress conditions, and the fundamental strategy can be adapted to investigate other species. Thus, TRIP is a highly complementary strategy to transposon-based approaches.

We first applied TRIP to MTB log-phase growth *in vitro* to characterize network perturbations that alter baseline fitness. **Figure 2A** visualizes abundance fold change of each TFI strain when induced in these conditions (see tab 1 of **Table S1** for detailed results). Most TFI strains showed no significant difference in abundance upon induction. Twenty-two TFI strains (10.6%; below dotted line at -0.5) exhibited substantial growth defects upon TF induction. (see Methods for details).

Growth-defective strains are enriched in TFs that activate genes associated with starvation responses (**Table S2**). Such strains are also enriched for TFs that repress essential genes (p < 10^−6^, hypergeometric test), although two defect-inducing TFIs (Rv3765c and Rv1255c) do not repress any essential genes, and 20 TFIs with no discernable growth phenotype do repress essential genes^7^. There is no significant correlation between the number of repressed essential genes and extent of growth defect incurred by the TFI strain.

To validate differences in relative abundance detected by TRIP, we compared the TRIP output with the growth of each individual TFI strain over a one-week time course with and without TF induction. Of the 22 TFI strains with strong growth defects in the TRIP assay, 20 also had strong growth defects when cultured individually. This validation indicates that 1) phenotypes detected by TRIP reflect growth observed in monoculture, and 2) significant growth defect upon TF induction is an uncommon and highly TF-specific effect.

Once baseline MTB network phenotypes were established, we applied TRIP to study response to INH. We exposed TFI pools to a dose of INH where the bulk population grew suboptimally (19% of untreated, see **Figure S1**), enabling us to identify TFIs with reduced and improved viability compared to the population average in a single experiment. **Figure 2D** shows abundance of each TFI strain exposed to drug in the induced condition relative to uninduced. TFI strains with a growth phenotype specific to INH partition into three groups: A) TFIs that convey growth advantage in INH but no change in when untreated (7 strains, purple box); B) TFIs that convey growth defect in INH but no change when untreated (4 strains, blue box); and C) TFIs that convey growth defect in both INH and untreated conditions (9 strains; light blue box). The genes whose promoters are bound by TFs in group A are enriched for transporter functions **Table S2**). In contrast, genes activated by TFs in groups B and C are significantly enriched for several stress response, homeostasis, and transport processes, and genes repressed by these TFs are significantly enriched for NADH dehydrogenase (ubiquinone) activity and DNA replication initiation (**Table S2**). Out of the 20 TFs associated with INH phenotypes by TRIP, two of them were reported by Xu and colleagues^16^ to alter MTB fitness significantly during INH treatment by Tn-seq (**Table S2**). The regulons of TFs in all three groups were enriched for genes reported to alter INH fitness significantly by Tn-seq (**Table S3**)^16^. Notably, two of the four TFs in group B solely activate genes when induced (Rv0330c and Rv2282c), so the association between members of these regulons and INH fitness could not have been detected by gene disruption-based assays.

The TFI conveying the greatest survival advantage during INH exposure is Rv1909c/ *furA*. This TF represses expression of *katG* (Rv1908c), which encodes the catalase-peroxidase enzyme that converts the INH prodrug into its active form^17,18^. Inducing *furA* in the presence of INH is known to restore nearly uninhibited growth^14,17,19^.

We next sought to investigate regulons that represented potential novel therapeutic targets. The TF conveying the greatest growth defect upon induction in INH is Rv1963c (*mce3R*), a TetR-like regulator. *mce3R* has previously been linked to the expression of genes that mediate β-oxidation of fatty acids and lipid transport^20–22^ and had no previous connection to INH.

To validate its hypersusceptibility, we tested viability of the *mce3R* induction strain in monoculture under INH treatment. First, we exposed the strain to INH, with and without induction of *mce3R* (**Figure 2E**). We confirmed induction of *mce3R* by qPCR (**Figure S2**), and observed a significant, 4-fold additional growth defect by 7 days of INH exposure, and a 21-fold additional defect by day 14. In addition, we also assayed MTB ATP levels (BacTiter Glo, Promega, Madison, WI) after 7 days of treatment with varying doses of INH (**Figure 2F**). We found that at every non-zero INH concentration we tested, induction of *mce3R* resulted in significantly lower metabolic viability, demonstrating that *mce3R*-mediated hypersusceptibility is independent of drug concentration.

**Figure 2.**
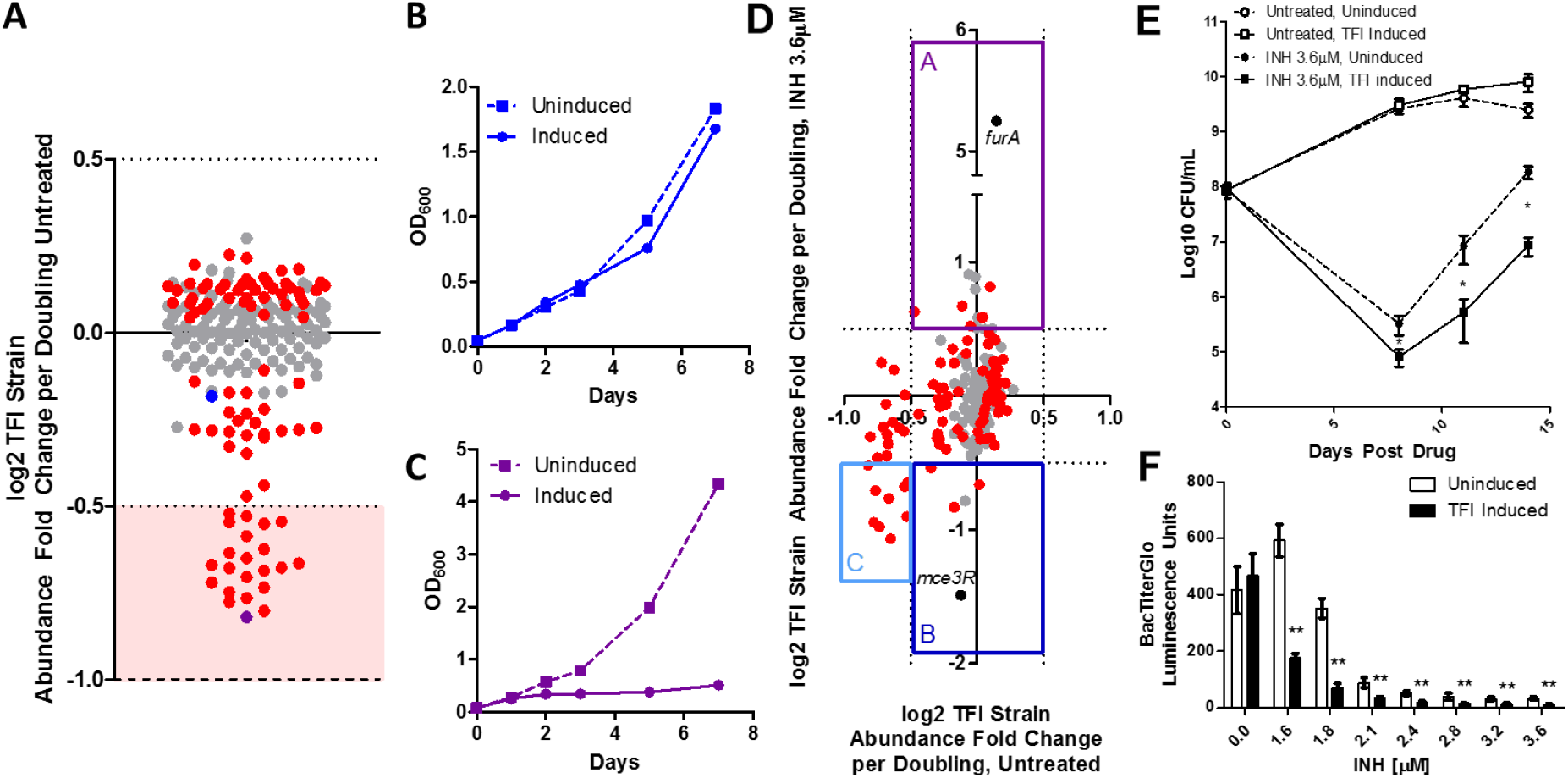
Regulon-mediated growth responses. (A) Log-phase TRIP results. Dots represent abundance change of TFI strains induced vs. uninduced, normalized by the estimated number of pool doublings. Red dots indicate z-score > 1, from three replicates. Dotted lines specify ± 2 standard deviations, and dashed line denotes detection limit, signifying no growth. Shaded area represents strains with strong defects. Monoculture growth curves of two strains (blue, purple dots) are shown in (B) and (C) respectively, with induction (solid) or without (dashed). (D) TRIP results with INH (y-axis) vs. no drug (x-axis). Since INH can be bactericidal, some strains showed abundance changes < –1. Boxes demarcate strains with altered INH survival. Black dots represent strains: Rv1909c (*furA*_TFI_) and Rv1963c (*mce3R*_TFI_). (E) *mce3R*_TFI_ colony forming units (CFU)/mL over 14 days in INH (black) vs. no drug (white) with induction (solid) or without (dashed). Error bars show standard deviation from three replicates. (F) *mce3R*_TFI_ metabolic viability after 7 days INH with induction (black) or without (white), measured by luminescence (see Methods). Error bars show standard deviation from four replicates. * and ** indicate significant differences between induction states (p < 0.05 and p < 0.001, Student’s t-test, respectively).

Hypersusceptibility could stem from some TFI-mediated countering of the MTB adaptation to INH. To investigate this hypothesis, we compared the *mce3R* induction regulon determined in our basal transcriptional network with the genes differentially expressed when H37Rv is exposed to INH^14,23,24^. Rv1469 is one of two genes repressed by *mce3R*(**Figure 3A**, see **Figure S3** for full set), and Rv1469 is normally upregulated in response to INH in broth culture and under macrophage infection conditions^23,24^. After ruling out the other gene (see **Table S4** and **Table S5** for details), we hypothesized that Rv1469 induction might be important for temporary MTB adaptation to INH. If so, reducing Rv1469 expression might contribute to INH hypersusceptiblity independent of expression changes in *mce3R*.

To test whether Rv1469 could influence INH susceptibility, we obtained a transposon mutant that disrupted Rv1469 (see Methods for details). We compared kill curves for this mutant (Rv1469::Himar1) vs. the parent strain CDC1551 when exposed to INH (**Figure 3B**). As predicted, survival was reduced in Rv1469::Himar1 relative to CDC1551 following INH treatment (42-fold difference after 7 days, p < 10^−4^, Student’s t-test), with no significant growth difference in absence of drug. Intriguingly, Rv1469 was not associated with significantly altered INH susceptibility in the Tn-seq screen by Xu and colleagues^16^.

**Figure 3.**
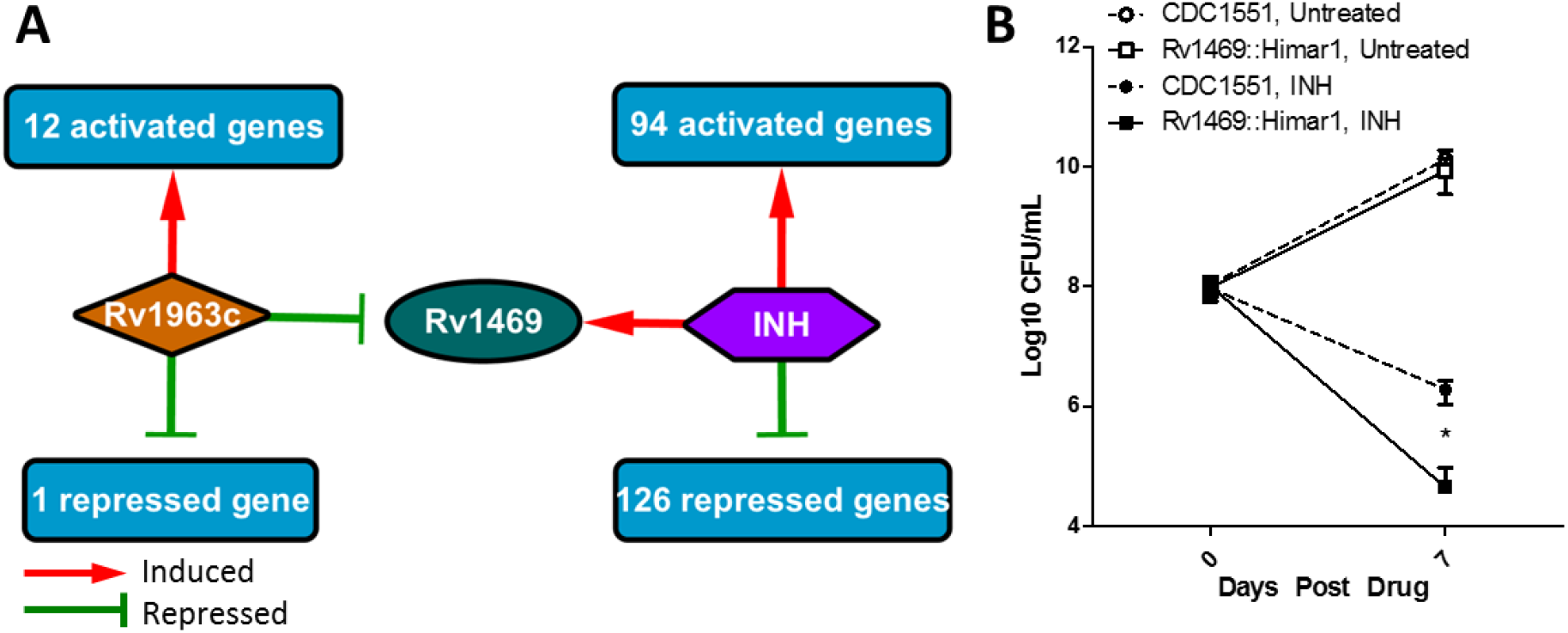
*mce3R* regulon reveals novel effector of INH susceptibility. (A) Network representation of overlap between *mce3R* regulon and genes differentially expressed in wildtype response to INH exposure. Red arrows indicate genes activated at least 2-fold, and green lines indicate genes repressed at least 2-fold. (B) CFU/mL over 7 days of Rv1469::Himar1 transposon disruption strain (solid) compared to the wildtype strain, CDC1551 (dashed). Both strains were exposed to 3.6µM INH (black) vs. no drug (white). Error bars indicate standard deviation from three replicates. The * indicates significant difference between the mutant and wildtype strains.

Rv1469 encodes a membrane protein^25^ annotated as the MTB homolog of CtpD, a member of the metal cation-transporting P1B4-ATPase subgroup, which is essential for MTB survival in the host^6,26,27^. CtpD is a high-affinity Fe^2+^ exporter needed for overcoming redox stress and adapting to the host environment^26,28^. Given that KatG-mediated catalysis is iron-dependent, Fe^2+^ accumulation resulting from loss of *ctpD* could possibly increase levels of oxy-ferrous KatG, which in turn could increase INH oxidation and activation^29^. Alternatively, extra free iron in MTB could promote cell wall changes^16^ or increased oxidative stress^30^ that may enhance INH activity. Further work is needed to establish the mechanism of CtpD-mediated INH tolerance, and whether this mechanism extends to other cation transmembrane transporters.

TRIP represents a powerful new tool to untangle the links between genetic perturbations and their phenotypic outcomes under specific environmental contexts. It can reveal novel associations between genes, networks and phenotype in several ways. First, by targeting regulons, TRIP harnesses nature’s levers to modulate responses— tuning gene sets that evolved to change coordinately—and uncovers emergent phenotypes that depend on synchronized actions of multiple genes. For example, two TFIs that slowed growth under log-phase conditions (Rv3765c and Rv1255c) do not repress any essential genes, suggesting that epistatic mechanisms may underlie these growth defects. Epistasis may also explain how TRIP successfully identified *mce3R* as a key regulatory potentiator of INH activity, whereas one of its target effector genes, *ctpD*, was not identified in a recent Tn-seq screen for INH-associated genes. In addition, TRIP samples network states that are distinct from those resulting from TF disruption. For example, *mce3R* was previously reported to regulate the *mce3* operon genes based on studies of a deletion mutant^20,21^. However, the transcriptional impact of inducing *mce3R* does not include the *mce3* operon (see tab 3 of Table S2 for full regulon, based on data from^14^), suggesting that *mce3R* participates in complex regulatory circuits. Combining gene disruption studies with TRIP screens and network analysis and could facilitate untangling these complex, nonlinear phenotypes. Finally, unlike gene-disruption assays, TRIP can identify fitness phenotypes associated with gene upregulation, such as in the cases of the INH hypersusceptibility-inducing TFs Rv0330c and Rv2282c, both of which exclusively activate genes when induced.

In the current study, we combined TRIP results with network analysis to identify novel genes that altered MTB response to the frontline drug INH. However, TRIP can interrogate network mediators of growth and survival under any condition from which microbes can be recovered, and TRIP requires tracking a substantially reduced set of mutants compared to Tn-seq, rendering it technically tractable. By integrating with network insight, TRIP will lend novel insights into emergent mechanisms underlying condition-specific growth phenotypes for MTB, and the strategy can be generalized for other organisms as well.

## Methods

### Strains and expression vectors

The individual strains comprising the MTB Transcription Factor Induction (TFI) Library were generated previously^14^. Briefly, 207 MTB DNA binding genes were cloned into a tagged, inducible vector that placed the TF under control of a tetracycline-inducible promoter and added a C-terminal FLAG epitope tag^14,31–34^. The constructs were then individually transformed into MTB H37Rv using standard methods. Individual TFI strains are available from the BEI strain repository at ATCC (^35^, NR-46512). The TFI library was generated by combining equal proportions of each strain into a common pool.

The Rv1469 transposon strain was obtained through BEI Resources, NIAID, NIH: MTB: Strain CDC1551, Transposon Mutant 1738 (MT1515, Rv1469), NR-18218^35^. The transposon insertion is located at base 671 in the 1974 base-pair long gene^35^.

### Culture

Bacteria were cultured at 37°C under aerobic conditions with constant agitation. For the experiments involving TFI strains, the strains were cultured in Middlebrook 7H9 with the ADC supplement (Difco), 0.05% Tween80, and 50 μg/mL hygromycin B to maintain the plasmids.

For the TRIP experiments, growth of the pooled TFI library was monitored by OD600. At an OD600 of 0.1, expression of the pooled TFI library was induced with anhydrotetracycline (ATc, 100ng/mL), and the cultures were grown for 7 days supplemented with either 3.6µM INH in 1% DMSO solution or DMSO as no-drug control. The cultures were sampled at Day 0 and Day 7 of the experiment for DNA isolation and subsequent sequencing.

For individual TFI strain time course experiments, each strain was cultured under the same media conditions as described for the pooled TFI library. When cultures reached OD600 ∼0.1, TFI strain induction and drug exposure proceeded as described for the pooled TFI library. The individual strain cultures were monitored for up to 14 days, with viability under the different drug and induction conditions assayed by plating on Middlebrook 7H10 solid media plates and assessing colony forming units using standard methods.

The Rv1469 transposon mutant strain was cultured in Middlebrook 7H9 with ADC supplement (Difco), 0.05% Tween80, and 30 μg/mL kanamycin to maintain the transposon insertion. Growth and survival of Rv1469 mutant was compared against the parent MTB CDC1551 strain. When cultures reached an OD600 of 0.1, drug exposure proceeded as described for the pooled TFI library. The individual strain cultures were monitored for up to 14 days, with viability under the different drug and induction conditions assayed by plating on Middlebrook 7H10 solid media plates and assessing colony forming units using standard methods.

### Dose-dependent viability assay

Strains were grown to log phase (OD600 ∼0.3), diluted to a final OD600 of 0.005, and dispensed into 96-well flat-bottom plates (Corning, Acton, MA) at a final volume of 200µL, containing 1% DMSO and varying concentrations of INH in the different wells. On each plate, control wells for each of the strains studied were included, containing no drug and 1% DMSO vehicle, to measure viability in the absence of INH exposure. Plates were incubated at 37°C for 7 days. Cellular viability was assayed on Day 7 by adding 20µL of culture from each well to 20µL of BacTiter-Glo Microbial Cell Viability Assay Reagent (Promega, Madison, WI), incubating at room temperature protected from direct light for 20 minutes, and reading luminescence intensity using a FluoStar Omega plate reader (BMG Lab Tech, Cary, NC).

### DNA isolation and sequencing

Cell pellets collected from each sample were resuspended in TE buffer, pH 8.0, transferred to a tube containing Lysing Matrix B (QBiogene, Inc.), and vigorously shaken three times at 6m/s for 30 seconds per cycle in a Bead Ruptor 24 homogenizer (Omni International, Kennesaw, GA), with a 30-second pause between each cycle. The mixture was centrifuged at maximum speed for one minute, and DNA was extracted from the supernatant using the MagJet Genomic DNA Kit (Thermo Fisher), according to the manufacturer’s instructions for manual genomic DNA purification.

PCR pre-amplification of DNA barcodes unique to each TFI strain was performed. The products of this reaction were prepared for Illumina sequencing using the NEBNext Ultra DNA Library Prep Kit for Illumina (New England Biolabs, Ipswich, MA) according to manufacturer’s instructions, and using the AMPure XP reagent (Agencourt Bioscience Corporation, Beverly, MA) for size selection and cleanup of adaptor-ligated DNA. We used the NEBNext Multiplex Oligos for Illumina (Dual Index Primers Set 1) to barcode the DNA libraries associated with each replicate and enable multiplexing of 96 libraries per sequencing run. The prepared libraries were quantified using the Kapa qPCR quantification kit, and were sequenced at the University of Washington Northwest Genomics Center with the Illumina NextSeq 500 Mid Output v2 Kit (Illumina Inc, San Diego, CA). The sequencing generated an average of 1.5 million 75 base-pair paired-end raw read counts per library.

### Sequencing read alignment and TFI strain abundance deconvolution

Read alignment was carried out using a custom processing pipeline that harnesses the Bowtie 2 utilities^36,37^, which is available at https://github.com/DavidRShermanLab/TRIPscreen, https://github.com/sturkarslan/DuffyNGS, and https://github.com/sturkarslan/DuffyTools. A custom Bowtie 2 target index was constructed from: the CDS sequences of all H37Rv genes; the inducible TFI anchor plasmid sequence; and the complete sequence of the empty plasmid as a negative control. The two mate-pair FASTQ datasets for each sample were separately mapped as unpaired reads using Bowtie 2’s local alignment mode. After both mate end datasets were aligned separately, the alignment results were combined to give a pair of gene/plasmid alignments for each raw read. Only raw read pairs having one alignment to the anchor plasmid and the other to a gene with an existing “Rv” code were kept as valid reads. Read pairs that mapped to “Rv” code genes on both ends, or pairs that failed to align were discarded. On average, each sample had 99.9% valid anchor/gene reads, which is comparable to typical RNA-seq and WGS alignment results. Libraries that generated fewer than 10,000 valid read pairs were excluded from further analysis. Valid reads were then tallied for all “Rv” code genes reported as raw abundance measures. Read counts for each TFI were then normalized as log2 reads per million (RPM) values. Higher RPM values indicated that the corresponding TFI strain had greater relative abundance in the pooled culture. The average log2 RPM values across TFIs were 11.7±3.2. TFIs with low abundance levels on day 0 of each experiment (log2 RPM < 5) were excluded from subsequent analysis (10 TFI strains, 4.8%).

To assess the effect of induction on TFI strain relative abundance, the log2 fold-change RPM values were calculated for the TFI-induced condition relative to un-induced. These values were further normalized by the number of doublings of the pooled library estimated from the change in OD600 over the course of the experiment. Positive fold change RPM values indicated that TFI induction conveyed a growth benefit, whereas negative fold change RPM values indicated that the TFI induction conveyed a growth defect under the conditions assayed. TFI strains that exhibited a log2 fold-change per doubling greater than 0.5 with z-score greater that 1 were deemed to have a significant growth phenotype under the condition assayed. The code for this processing is available at https://github.com/DavidRShermanLab/TRIPscreen.

### Statistics

Unless indicated, experiments were performed three times, and the mean and standard deviation from biological replicates of representative experiments are reported. Statistical differences between means were evaluated by two-tailed Student’s t-tests, statistically significant correlation was evaluated by calculating a Pearson correlation coefficient and comparing against a Student’s t distribution, and statistical enrichment was evaluated by hypergeometric test, unless otherwise noted. The significance cutoff was set at p < 0.05, unless otherwise indicated.

### Gene Ontology Enrichment Analysis

The gene ontology (GO) term annotations for genes comprising the regulons of the TFs under analysis were taken from^38^ and evaluated for statistical enrichment against the GO annotations for the entire gene set of the MTB strain H37Rv using the hypergeometric test and further subjected to a Bonferroni correction for multiple hypothesis testing, with the number of independent tests estimated as the number of GO terms associated to at least 2 genes in the H37Rv reference gene set (analogous to method used by^39^). We further filtered the enriched GO terms to only those featured in the regulons for 2 more of the TFs under analysis.

### Data Availability

The data reported in the paper are available in the Supplementary Materials. The raw. fastq sequence data files are deposited in the Sequence Read Archive database [accession number pending]. The code required to process the sequenced reads are available at: https://github.com/sturkarslan/DuffyNGS, https://github.com/sturkarslan/DuffyTools, and https://github.com/DavidRShermanLab/TRIPscreen.

## Acknowledgments

We gratefully acknowledge Jessica Winkler, Jenny Lohmiller, and Reiling Liao for their technical assistance, and we also thank Kyu Rhee and Susan Shen for helpful discussions.

This work was supported by the National Institutes of Health [grant numbers U 19 AI106761; U19 AI111276; U19 AI135976; 5T32AI007509].

## Author Contributions

S.M. conceived of the study, led the design, generated data, analyzed the results, and drafted the manuscript. B.M. developed the software to convert raw sequencing data into abundance values for each TFI strain. S.H. generated data and assembled the pooled TFI library cultures. J.F. assisted with sample preparation for sequencing. T.R. and D.S. conceived of the study, led the design, organized the data analysis, and drafted the manuscript.

## Materials & Correspondence

Correspondence and material requests should be addressed to: David Sherman.

## Supplementary Information

**Figure S1.**
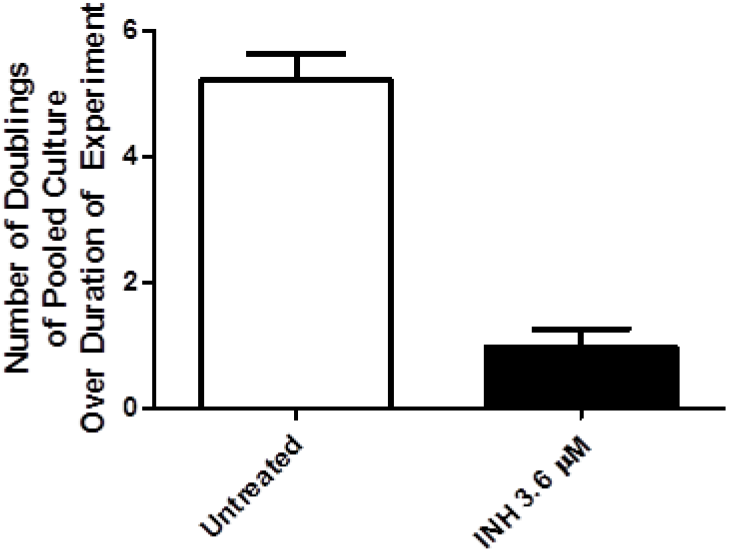
Comparing TFI pool growth between experimental conditions. Number of doublings for TFI strain pool over duration of TRIP experiments in the untreated vs. INH treated conditions, estimated from the change of OD600 over the course of the experiment.

**Figure S2.**
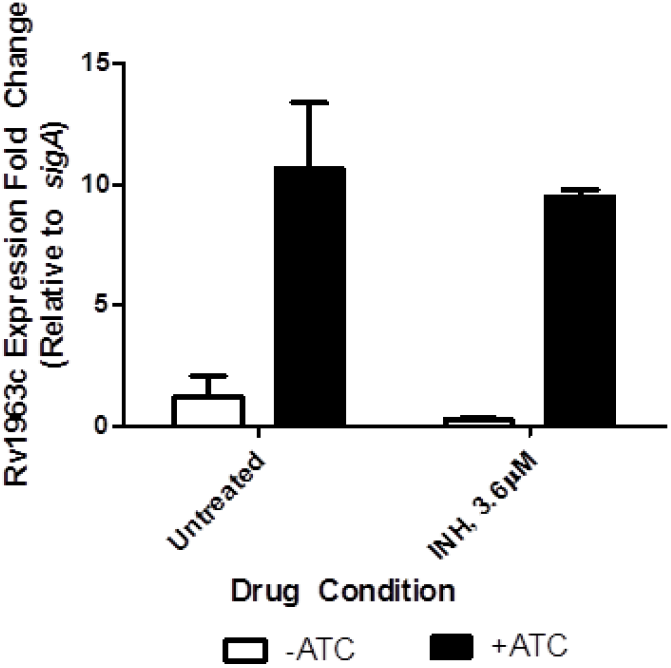
Chemical induction triggers *mce3R* expression change. Expression fold change of Rv1963c relative to the housekeeping gene sigA, assessed by qPCR, representing 3 biological replicates from a representative experiment (two were performed in total). Conditions compared are in absence (white bars) and presence (black bars) of anhydrous-tetracycline (ATc) inducer, and presence and absence of INH exposure. Results show at least 8-fold activation of Rv1963c expression upon induction with ATC in both absence and presence of INH (p < 0.05, Student’s t-test).

**Figure S3.**
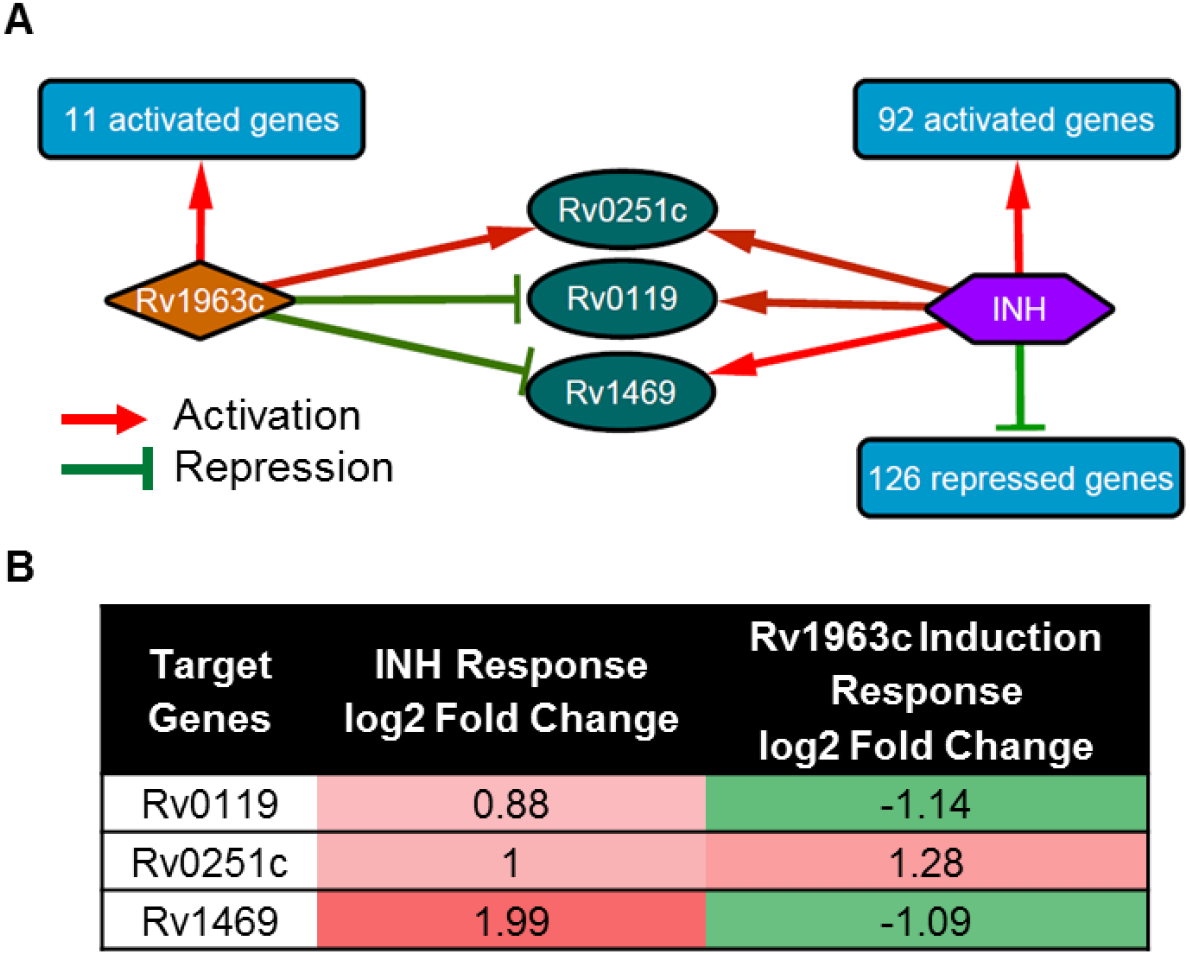
Overlap of genes regulated by *mce3R* that also modulate expression in baseline response to INH exposure. (**A**) Network diagram depicts the genes differentially expressed upon induction of Rv1963c/*mce3R* expression (left), and upon exposure to INH (right). Three genes alter expression under both these conditions. (**B**) Table summarizes the expression fold-changes of the genes perturbed both by *mce3R* induction and INH exposure.

**Table S1.** (See TableS1.xlsx). Detailed results of TRIP screens in presence and absence of 3.6µM INH. Tab 1 reports mean log2 abundance fold change for each TFI strain, relative to no induction. Tab 2 reports the log2 RPM values for each sample analyzed for the study.

**Table S2.** (See TableS2.xlsx). Detailed regulon and accompanying GO term enrichment information of TFs that convey altered fitness under different conditions. GO annotations were from^38^. Tab 1 provides information for TFs conveying growth phenotype under log-phase growth conditions. Tab 2 provides information for TFs conveying conditional resistance to INH (Group A). Tab 3 provides information for TFs conveying hypersusceptibility to INH but no growth defect in absence of drug (Group B). Tab 4 provides information for TFs conveying hypersusceptibility to INH and growth defect in absence of drug (Group C). Tab 5 provides information for all TFs conveying hypersusceptibility to INH (Groups B and C combined). The regulon information are based on data generated from^13,14^. The essentiality information is based on data generated from^7,16^.

**Table S3.** (See TableS3.xlsx). Overlap of TFI regulons with INH growth phenotype and genes with INH fitness phenotype in Tn-seq assay measured by^16^.

**Table S4.** Correlation between Rv1469 regulation and INH response phenotype. Summary of regulatory impact and TRIP abundance fold change upon INH exposure for the TFs that regulate Rv1469. Note that Rv1990c and Rv0022c do not have TRIP values associated because the raw sequence data had poor detection of these TFI strains under all conditions. Expression fold change values were assembled from^14^.

| TF | Gene Name | Total Genes Regulated | Rv1469 Expression Fold Change upon TF Induction (No Drug) | TRIP Abundance Fold Change, INH |
| --- | --- | --- | --- | --- |
| Rv3852 | *hns* | 7 | 1.05 | 0.42 |
| Rv1963c | *mce3R* | 14 | -1.09 | -1.48 |
| Rv1221 | *sigE* | 25 | -1.10 | -0.28 |
| Rv1049 | *mosR* | 74 | -1.54 | -0.29 |
| Rv1990c |  | 91 | -1.27 | NA |
| Rv0757 | *phoP* | 139 | -1.08 | -0.32 |
| Rv0324 |  | 254 | -1.21 | -0.67 |
| Rv0238 |  | 353 | -1.09 | 0.09 |
| Rv3223c | *sigH* | 359 | -1.77 | -0.53 |
| Rv3416 | *whiB3* | 371 | -1.96 | -0.64 |
| Rv0022c | *whiB5* | 435 | -1.82 | NA |

**Table S5.** Correlation between Rv0119 regulation and INH response phenotype. Summary of regulatory impact and TRIP abundance fold change upon INH exposure for the TFs that regulate Rv0119, the other gene induced during basal INH exposure in H37Rv and repressed by *mce3R*.

| TF | Gene Name | Total Genes Regulated | Rv0119 Expression Fold Change upon TF induction (No Drug) | TRIP Abundance Fold Change, INH |
| --- | --- | --- | --- | --- |
| Rv0117 | *oxyS* | 4 | -1.58 | -0.11 |
| Rv3055 |  | 6 | -1.24 | 0.07 |
| Rv1963c | *mce3R* | 14 | -1.14 | -1.48 |
| Rv0260c |  | 235 | -1.43 | 0.04 |
| Rv0324 |  | 254 | 1.03 | -0.67 |
| Rv3223c | *sigH* | 359 | 1.28 | -0.53 |
| Rv0081 |  | 402 | 1.95 | -0.76 |
| Rv0023 |  | 893 | 1.17 | -1.05 |

